# Inorganic Nitrogen Availability Drives Metabolic Specialization and Adaptive Strategies in *Vibrio harveyi* and *Vibrio parahaemolyticus*

**DOI:** 10.64898/2026.08.10.743883

**Authors:** Xi Xiong, Hui Ren, Shijun Chen, Lian Gan

## Abstract

Nitrogen availability is a key factor shaping microbial metabolism, ecological adaptation, and nitrogen cycling in aquatic environments. Members of the genus *Vibrio* are ubiquitous heterotrophic bacteria in marine and aquaculture ecosystems, yet their responses to different inorganic nitrogen sources remain poorly understood. Here, we systematically compared the growth characteristics, nitrogen transformation capacity, and molecular responses of *Vibrio harveyi* and *Vibrio parahaemolyticus* under ammonium (NH_4_^+^), nitrate (NO_3_^−^), and nitrite (NO_2_^−^) conditions using physiological assays, comparative genomic analysis, and transcriptomic profiling. *V. harveyi* exhibited broader nitrogen utilization capacity and was able to grow under all three nitrogen conditions, whereas *V. parahaemolyticus* showed a strong preference for NH_4_^+^ and limited growth under NO_3_^−^ and NO_2_^−^ conditions. Moreover, *V. harveyi* displayed rapid population expansion accompanied by reduced long-term viability, while *V. parahaemolyticus* maintained greater population stability. Both species showed NO_3_^−^ accumulation during growth despite lacking canonical nitrification genes under NH_4_^+^ condition, suggesting the potential involvement of non-canonical heterotrophic nitrification processes. Transcriptomic analysis revealed nitrogen source-dependent metabolic specialization in *V. harveyi*. NH_4_^+^ availability promoted motility-associated responses and metabolic overflow, whereas NO_3_^−^ induced iron acquisition-related pathways and NO_2_^−^ activated assimilatory nitrite reduction coupled with oxidative stress adaptation. These findings demonstrate that inorganic nitrogen availability drives divergent metabolic and adaptive strategies in *Vibrio*, providing new insights into their nitrogen metabolic potential and ecological roles in aquatic environments.

**Importance:** This study demonstrates that *V. harveyi* and *V. parahaemolyticus* exhibit distinct inorganic nitrogen utilization strategies, with *V. harveyi* displaying broader nitrogen utilization capacity. Transcriptomic and metabolomic analyses revealed that different nitrogen sources drive distinct metabolic and environmental adaptation responses in *V. harveyi*, including enhanced motility-associated functions and metabolic overflow responses under NH_4_^+^ condition, increased iron acquisition pathways under NO_3_^−^ condition, and activation of assimilatory nitrite reduction coupled with oxidative stress adaptation under NO_2_^−^ condition. Furthermore, significant nitrate accumulation was observed in both Vibrio strains during ammonium cultivation despite the absence of canonical nitrification genes, suggesting unexplored nitrogen transformation potential in vibrios. This study expands our understanding of how inorganic nitrogen availability shapes microbial adaptation strategies and ecological functions in aquatic environments.

## Introduction

Nitrogen availability is a major environmental factor shaping microbial metabolism, ecological interactions, and biogeochemical cycling in aquatic ecosystems^1^. Microorganisms regulate nitrogen fluxes through a variety of classical nitrogen metabolic pathways, including ammonium assimilation, nitrate and nitrite assimilation, dissimilatory nitrate reduction to ammonium (DNRA), denitrification, urea utilization, and other nitrogen transformation processes^1–11^. These processes collectively drive the transformation of nitrogen compounds and regulate their availability in both aquatic and terrestrial environments^12,13^. Among inorganic nitrogen forms, ammonium (NH_4_^+^), nitrate (NO_3_^−^), and nitrite (NO_2_^−^) serve not only as essential nitrogen sources for microbial biosynthesis but also as important environmental factors that impose distinct metabolic constraints and selective pressures^14,15^. Therefore, understanding how microorganisms respond to different inorganic nitrogen sources is critical for elucidating microbial adaptation strategies and their roles in aquatic nitrogen cycling.

Members of the genus *Vibrio* are widespread heterotrophic bacteria in marine, estuarine, and aquaculture ecosystems, where they contribute to organic matter degradation, nutrient transformation, and microbial community dynamics. Their rapid growth, metabolic flexibility, and ability to respond to environmental fluctuations enable them to occupy diverse ecological niches, including seawater, sediments, and host-associated environments^16^. Meanwhile, several *Vibrio* species are important opportunistic pathogens of aquatic animals and humans, and their environmental persistence and host colonization capacity are closely associated with their adaptive metabolic strategies^17^. Previous genomic studies have revealed that *Vibrio* species harbor diverse nitrogen metabolism-related genes, indicating their broad potential for inorganic nitrogen acquisition and transformation. Meanwhile, nitrogen metabolism-related gene repertoires vary substantially across *Vibrio*, reflecting divergent nitrogen metabolic capacities. For example, *V. cholerae* lacks canonical denitrification-associated nitrite reductase genes^18^. Moreover, *Vibrio* genomes generally lack canonical denitrification-associated nitrite reductases (*nirS* and *nirK*), while often encoding the assimilatory nitrite reductase system *nirBD*^9^.

Previous studies have shown that *Vibrio* utilize diverse nitrogen sources to support growth and ecological adaptation, with nitrogen metabolism also contributing to pathogenic regulation and marine nitrogen cycling^19–21^. However, whether inorganic nitrogen availability shapes metabolic specialization and adaptive strategies in Vibrio remains poorly understood. Different nitrogen sources impose distinct metabolic demands: NH_4_^+^ can be directly assimilated into biosynthetic pathways, whereas NO_3_^−^ and NO_2_^−^ require additional reduction steps prior to assimilation. Such differences may influence bacterial growth, metabolic regulation, and stress adaptation across environmental nitrogen regimes.

Here, we investigated the effects of different inorganic nitrogen sources on the growth characteristics, nitrogen transformation capacity, and adaptive responses of *Vibrio harveyi* and *Vibrio parahaemolyticus*. By integrating physiological assays, nitrogen transformation measurements, comparative genomic analysis, transcriptomics, and metabolomics, we aimed to elucidate how inorganic nitrogen availability regulates nitrogen utilization strategies and drives metabolic specialization in *Vibrio*. This study provides new insights into the relationship between environmental nitrogen availability, microbial adaptation, and ecological functions of *Vibrio* in aquatic ecosystems.

## Results

### Different nitrogen sources drive distinct population growth and viability maintenance strategies in *Vibrio*

Two bacterial isolates, *V. harveyi* Vh-1 and *V. parahaemolyticus* Vp-64, were isolated from crustaceans collected from farms in southern China. Both strains have been subjected to whole-genome sequencing and deposited in NCBI. To investigate differences in nitrogen source utilization between the two *Vibrio* strains, we optimized the M9 medium composition (see Methods). Briefly, glucose was provided as the sole carbon source, while sodium nitrate (NO_3_^−^), ammonium chloride (NH_4_^+^), and sodium nitrite (NO_2_^−^) were supplied individually as the sole nitrogen sources. To exclude the potential influence of residual nutrients from the LB activation culture, nitrogen-free medium was used as a control. After cultivation in nitrogen-free medium, neither strain formed colonies when plated onto thiosulfate–citrate–bile salts–sucrose (TCBS) agar, indicating that both strains were unable to maintain growth and proliferation in the absence of an external nitrogen source. Therefore, the bacterial growth observed in subsequent experiments was primarily supported by the inorganic nitrogen sources provided in the medium rather than residual nitrogen carried over from LB activation.

We first evaluated bacterial growth under different inorganic nitrogen conditions by measuring OD_600_. Analysis of slope changes in the growth curves revealed that, under NH_4_^+^ condition, both strains exhibited rapid biomass accumulation during the early cultivation period, with OD_600_ values increasing within 24 h and subsequently declining after prolonged incubation (168 h). Both strains were able to utilize NH_4_^+^ as a nitrogen source to support growth, while Vp-64 exhibited higher overall OD_600_ values and a larger area under the curve (AUC) than Vh-1, indicating a stronger growth capacity under NH_4_^+^ supplementation. (Fig. 1a, b).

**Figure 1.**
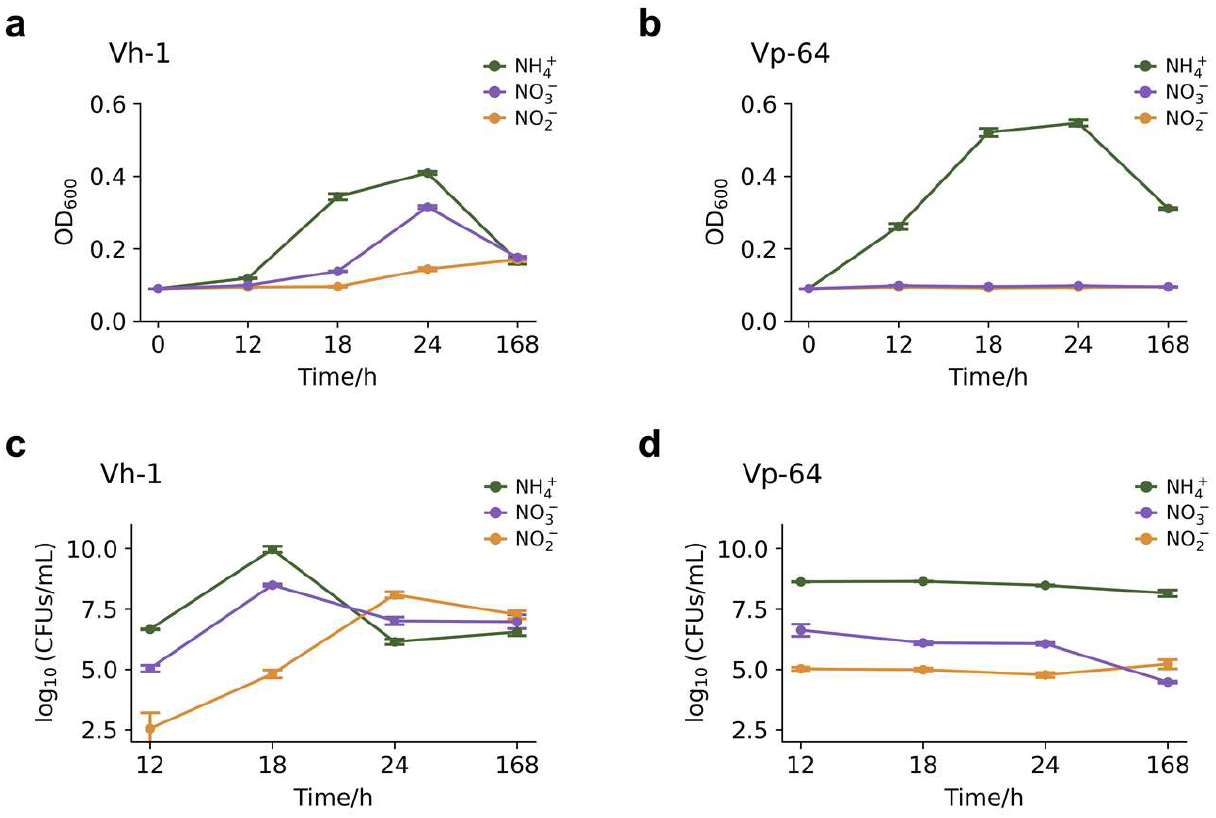
Growth performance and viability dynamics of two *Vibrio* strains. Error bars represent standard error of the mean (SEM), and viable cell counts are expressed as log_10_ CFUs/mL. **(a–b)** OD_600_-based growth curves of Vh-1 (a) and Vp-64 (b). **(c–d)** Viable cell count dynamics of Vh-1 (c) and Vp-64 (d), represented as log_10_ CFUs/mL.

However, the two strains displayed distinct growth patterns under NO_3_^−^ and NO_2_^−^ conditions. Vh-1 was still capable of growth under NO_3_^−^ and NO_2_^−^, although its growth rate and final biomass were lower than those observed under NH_4_^+^ condition. Among these conditions, NO_2_^−^ resulted in the smallest AUC, reflecting the lowest overall biomass accumulation, indicating that NO_2_^−^ was less favorable for supporting Vh-1 growth compared with other tested nitrogen sources. In contrast, Vp-64 maintained OD_600_ values close to the initial level (<0.1) throughout cultivation with NO_3_^−^ or NO_2_^−^, indicating that substantial biomass accumulation was not observed under these conditions. These results indicate that Vp-64 lacks the ability to effectively utilize NO_3_^−^ and NO_2_^−^ to support population expansion, whereas Vh-1 possesses a broader inorganic nitrogen utilization capacity.

Because OD_600_ reflects total biomass rather than viable bacterial numbers, we further quantified viable cells using TCBS plate counting to evaluate changes in population viability under different nitrogen conditions (Fig. 1c, d). By comparing the slope changes of the viability curves over the same time interval, we found that under NH_4_^+^ condition, although the OD_600_ of Vh-1 increased from 18 h to 24 h, the viable cell number decreased during the same period, suggesting that rapid biomass accumulation did not necessarily correspond to increased population viability. In contrast, Vp-64 maintained relatively stable viable cell numbers during rapid growth under NH_4_^+^ condition, indicating stronger population stability.

Further analysis of viable cell dynamics showed that Vh-1 grown with NO_3_^−^ exhibited a similar pattern to that observed under NH_4_^+^ condition, with viable cell numbers declining after rapid biomass accumulation. However, under NO_2_^−^ condition, viable cell numbers gradually increased along with biomass accumulation. For Vp-64, although NO_3_^−^ and NO_2_^−^ did not support significant population expansion, viable cell numbers remained relatively stable, further demonstrating its stronger ability to maintain population viability under unfavorable nitrogen conditions.

We additionally monitored changes in medium pH during cultivation. Vh-1, which was capable of growth under all three nitrogen conditions, exhibited a continuous decrease in medium pH, suggesting the accumulation of acidic metabolic products during growth. Similarly, Vp-64 showed a decrease in pH only under NH_4_^+^ condition where active growth occurred, whereas no significant pH changes were observed under NO_3_^−^ or NO_2_^−^ condition where population expansion was absent.

Collectively, these results demonstrate that different inorganic nitrogen sources drive distinct growth and survival strategies in *Vibrio*. Vh-1 exhibited broader nitrogen utilization capacity and supported population expansion using NH_4_^+^, NO_3_^−^, and NO_2_^−^. However, increased biomass accumulation was not always coupled with enhanced viability, with this discrepancy being particularly evident under NH_4_^+^ and NO_3_^−^ conditions. In contrast, Vp-64 primarily relied on NH_4_^+^ for population expansion. However, it maintained stronger long-term viability under alternative inorganic nitrogen conditions despite limited biomass accumulation. These findings suggest that the two *Vibrio* strains employ distinct ecological strategies in response to inorganic nitrogen availability.

### Dynamic changes in inorganic nitrogen reveal potential heterotrophic nitrification capacity in *V. harveyi* and *V. parahaemolyticus*

To further investigate nitrogen utilization strategies of the two *Vibrio* strains under nitrogen conditions, we monitored dynamic changes in three major inorganic nitrogen forms (NH_4_^+^, NO_2_^−^, and NO_3_^−^) during cultivation.

Under NH_4_^+^ condition, OD_600_ measurements and TCBS viable cell counts demonstrated active population expansion of both *Vibrio* strains. Correspondingly, NH_4_^+^ concentrations in the culture medium rapidly decreased and were almost depleted within 18 h, indicating that NH_4_^+^ was actively utilized to support bacterial growth (Fig.2a, b). Interestingly, substantial accumulation of NO_3_^−^ was detected during cultivation, suggesting that both strains possessed the capacity to convert NH_4_^+^ into NO_3_^−^ (Fig. 2c, d). These results indicated a potential heterotrophic nitrification process in *Vibrio*.

**Figure 2.**
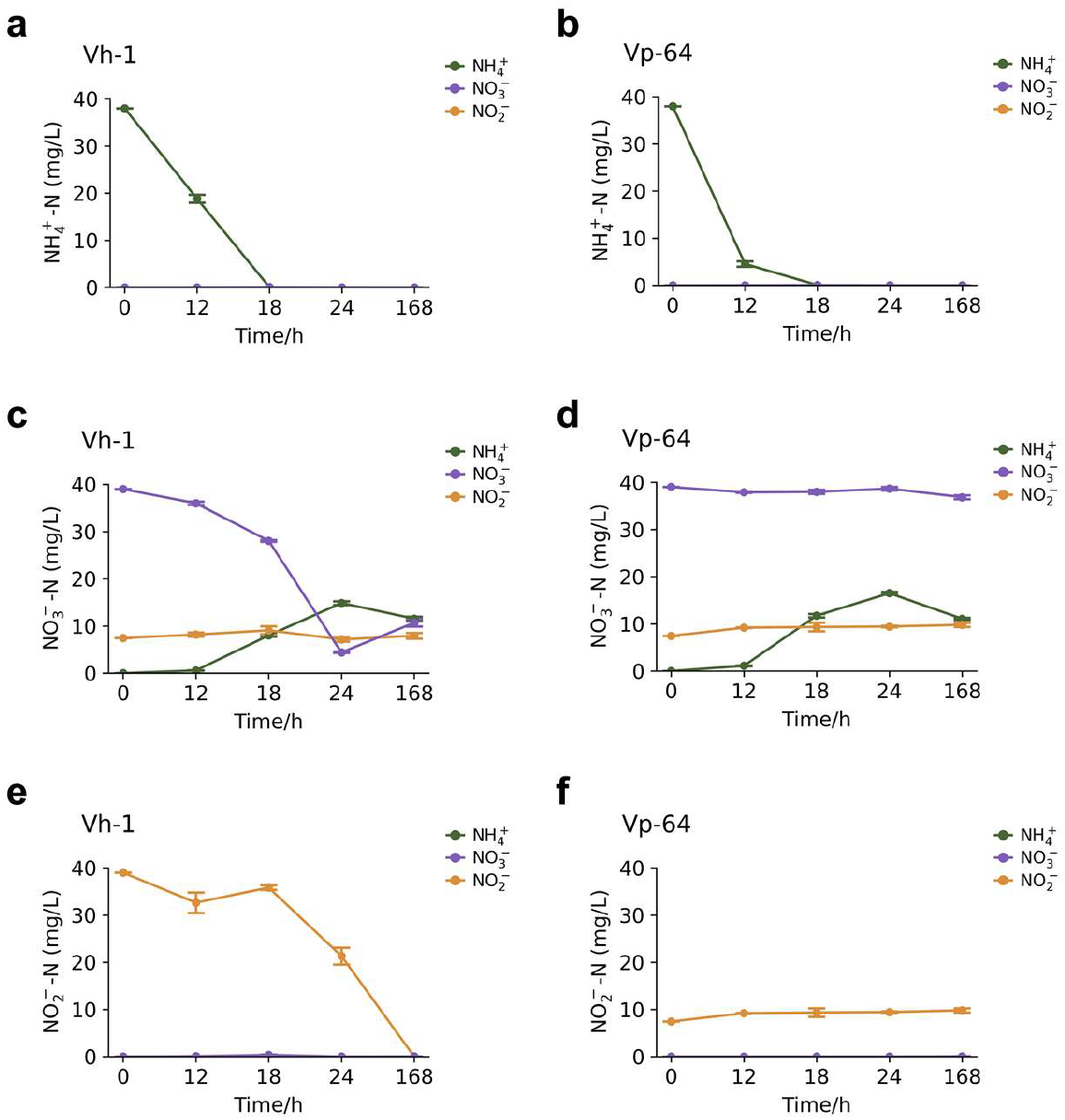
Dynamics of inorganic nitrogen concentrations in the culture media of two *Vibrio* strains. Nitrogen concentrations are expressed as elemental N mass concentrations. Error bars represent SEM, and color labels indicate the corresponding cultivation conditions. **(a–f)** Changes in NH_4_^+^-N, NO_3_^−^-N, and NO_2_^−^-N concentrations in Vh-1 **(a, c, e)** and Vp-64 **(b, d, f)** under different cultivation conditions.

For Vh-1, transient accumulation of trace NO_2_^−^ was observed under NH_4_^+^ condition at 12 h and 18 h, whereas NO_2_^−^ was no longer detectable at 24 h (Fig.2a). Under NO_2_^−^ condition, Vh-1 utilized NO_2_^−^ as a nitrogen source, and no residual NO_2_^−^ was detected in the culture medium after 168 h, demonstrating its ability to utilize nitrite for growth (Fig.2e). However, after NO_2_^−^ depletion, low levels of NO_3_^−^ were detected in the medium at 168 h, suggesting that NO_2_^−^ may have been partially converted to NO_3_^−^ by Vh-1 and raising the possibility of heterotrophic nitrification activity (Fig.2c).

For Vp-64, as described above, NO_3_^−^ continuously accumulated in the medium under NH_4_^+^ condition, without detectable NO_2_^−^ accumulation. When NO_3_^−^ was supplied as the sole nitrogen source, Vp-64 showed no apparent NO_3_^−^ consumption, with only trace NO_2_^−^ production detected after prolonged cultivation (168 h), consistent with its limited ability to utilize NO_3_^−^ for growth (Fig.2d, f). These results suggest distinct nitrogen transformation patterns between the two *Vibrio* species.

Classical nitrification involves the stepwise oxidation of ammonium to nitrate through ammonia oxidation and nitrite oxidation, mediated by specialized nitrifying microorganisms and key enzymes including ammonia monooxygenase (*amo*), hydroxylamine dehydrogenase (*hao*), and nitrite oxidoreductase (*nxr*)^5,6,10,11^. To investigate the potential genetic basis activity in both strain, we integrated RefSeq genome annotations with additional annotation using the NCycDB (S. Table 1)^22^. Genome annotation revealed that Vh-1 and Vp-64 possessed genes associated with multiple nitrogen metabolic pathways, including periplasmic nitrate reduction (*napABC*), assimilatory nitrate reduction (*nasA*), assimilatory nitrite reduction (*nirBD*), DNRA (*nrfABCD*), ammonium assimilation pathways (*glnA*, *gltBD*), and urea utilization (*ureABC*). These findings indicate that both strains possess extensive genetic potential for inorganic nitrogen transformation. However, neither strain contained the canonical nitrification genes involved in ammonia oxidation (*amoABC*, *hao*) or nitrite oxidation (*nxrAB*). Furthermore, analysis of approximately 30,000 publicly available *Vibrio* genomes from NCBI revealed a general absence of these canonical nitrification genes across the genus, suggesting that the classical nitrification pathway is unlikely to occur in *Vibrio*.

Given that nitrite oxidation represents the terminal step in nitrate formation, we performed protein structure prediction and structural similarity analysis against *nxrA* to identify potential molecular candidates involved in nitrate production. Several proteins belonging to the molybdenum enzyme superfamily, including *napA*, *torA*, *fdhA*, and *nasA*, exhibited high structural similarity to *nxrA* (TM-score >0.5, coverage >70%) based on structural comparisons (Supplementary Material). Previous biochemical studies have also suggested that *napA* may possess nitrite oxidation activity under in vitro conditions^23^. Pairwise sequence comparisons between *napA* and known *nxrA* proteins revealed relatively low sequence identities (<35%).

Collectively, these observations suggest that nitrate production in *Vibrio* is unlikely to be mediated by canonical *amo*, *hao*, or *nxrA* genes, but may instead involve alternative molybdenum-dependent enzymes or non-classical nitrogen transformation pathways.

### Nitrogen source-dependent regulation of nitrogen assimilation pathways in *V. harveyi*

Although Vh-1 exhibited a decline in viable cell numbers during early cultivation, the timing of biomass plateau formation varied substantially among different inorganic nitrogen sources. Under NH_4_^+^ condition, NH_4_^+^ was rapidly depleted during the early growth stage, and the OD_600_ plateau was reached at approximately 35 h. The plateau phase under NO_3_^−^ condition was established later, at approximately 43 h, whereas NO_2_^−^ condition showed the latest plateau formation at approximately 59 h. Therefore, bulk RNA-seq analysis was performed using samples collected at the biomass plateau stage to characterize the transcriptional responses and potential regulatory mechanisms underlying nitrogen metabolism in Vh-1 under different inorganic nitrogen conditions.

Principal component analysis (PCA) and hierarchical clustering demonstrated clear separation of transcriptomic profiles among the three nitrogen treatments, indicating that different nitrogen sources induced distinct transcriptional programs (Fig. 3a). A subset of randomly selected genes was further validated by qPCR, and the expression trends were consistent with RNA-seq results, confirming the reliability of the transcriptomic data (S. Fig. 1).

**Figure 3.**
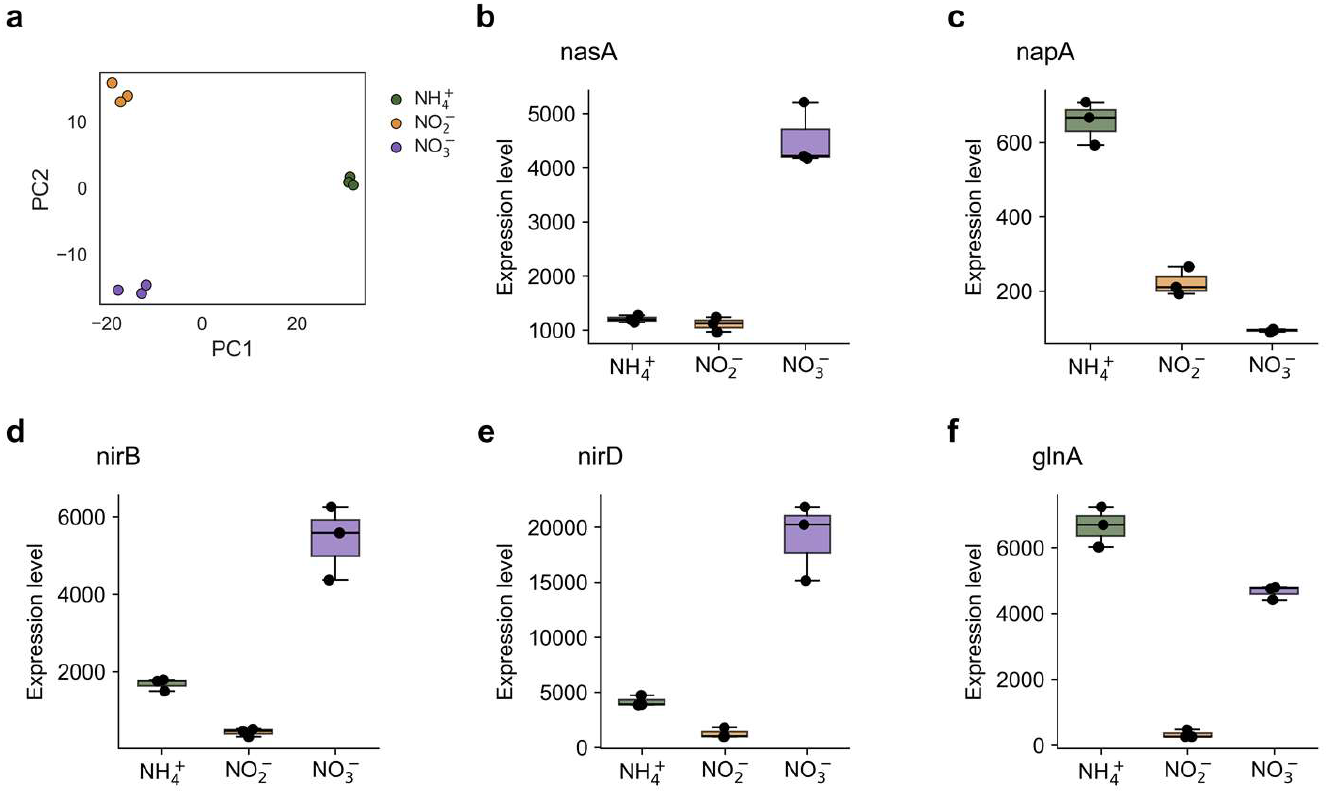
Expression profiles of nitrogen metabolism-related genes under different inorganic nitrogen conditions. **(a)** PCA of RNA-seq samples. **(b–f)** Relative expression levels of nitrogen metabolism-related genes, including *nasA*, *napA*, *nirB*, *nirD* and *glnA*, represented by DESeq2-normalized read counts.

We analyzed the expression patterns of nitrogen metabolism-related genes. Under NO_3_^−^ cultivation, the *nasA* exhibited significantly higher expression levels compared with NH_4_^+^ and NO_2_^−^ cultivation groups, indicating that *V. harveyi* induced an assimilatory nitrate reduction system under nitrate conditions (Fig. 3b). This system likely reduces nitrate to nitrite, which is subsequently incorporated into cellular nitrogen metabolism, thereby enabling *V. harveyi* to utilize nitrate as a nitrogen source for growth.

Interestingly, the periplasmic nitrate reductase gene *napA* showed significantly higher expression under NH_4_^+^ cultivation than under NO_3_^−^ cultivation (Fig. 3c). Combined with the observation that NH_4_^+^ cultivation resulted in nitrate accumulation, we speculate that *napA* may participate in a non-classical nitrogen transformation process under carbon-rich conditions, potentially contributing to nitrite-to-nitrate conversion. However, further isotope tracing and biochemical characterization are required to confirm this hypothesis.

The key gene *nrfA* involved in DNRA showed little or no detectable expression under all three nitrogen conditions. In contrast, the assimilatory nitrite reductase genes *nirB* and *nirD* exhibited clear expression patterns (Fig. 3d, e). These results suggest that nitrite utilization in *V. harveyi* mainly depends on assimilatory nitrite reduction rather than DNRA, where nitrite serves as a terminal electron acceptor for energy conservation. Interestingly, the *nirBD* gene cluster showed the highest expression under NO_3_^−^ condition, followed by NH_4_^+^ cultivation, whereas the lowest expression was observed under NO_2_^−^ condition. This pattern suggests that *nirBD* expression is not solely induced by nitrite availability but is also regulated by metabolic adaptation strategies associated with nitrite flux limitation or potential nitrite stress.

Meanwhile, the glutamine synthetase gene *glnA* exhibited high expression under both NH_4_^+^ and NO_3_^−^ conditions, indicating active incorporation of ammonium into glutamate metabolism and downstream organic nitrogen biosynthesis (Fig. 3f). Since no significant ammonium accumulation was detected in NH_4_^+^ cultures, this suggests that intracellular ammonium generated or absorbed by cells was rapidly assimilated. In contrast, *glnA* expression remained relatively low under NO_2_^−^ cultivation, which may reflect limited nitrogen flux and reduced nitrogen assimilation demand associated with slower growth under nitrite conditions.

### Ammonium promotes motility-associated transcriptional responses and induces metabolic overflow in *Vibrio*

Because Vh-1was able to produce nitrate under NH_4_^+^ cultivation and utilize both NH_4_^+^ and NO_3_^−^ for growth, we compared global transcriptional profiles between these two nitrogen conditions to investigate nitrogen source-dependent physiological adaptation. Differential expression analysis identified 456 genes significantly upregulated under NH_4_^+^ cultivation and 384 genes significantly upregulated under NO_3_^−^ cultivation. GO enrichment analysis revealed that both conditions shared conserved cellular processes, including DNA-templated transcription, DNA repair, and proteolysis (S. Fig. 2). However, NH_4_^+^ cultivation specifically induced genes associated with bacterial flagellar assembly. Consistently, the coordinated upregulation of key flagellar genes, such as the regulatory components *flrBC*, the flagellar export apparatus component *flhA*, the basal body protein *fliF*, and the motor protein gene *motA*, suggests an enhanced capacity for flagellar assembly and motility. In contrast, the relatively reduced expression of pilus- and capsular polysaccharide-associated genes (*msh, cps*) suggests a reduced investment in surface adhesion and extracellular matrix production (Fig. 4a). Weighted gene co-expression network analysis (WGCNA) further identified a NH_4_^+^-responsive MEblue module containing numerous flagellar-associated genes, which showed a strong positive correlation with NH_4_^+^ cultivation, suggesting enhanced motility and nutrient-searching capacity under ammonium availability (Fig. 4b).

**Figure 4.**
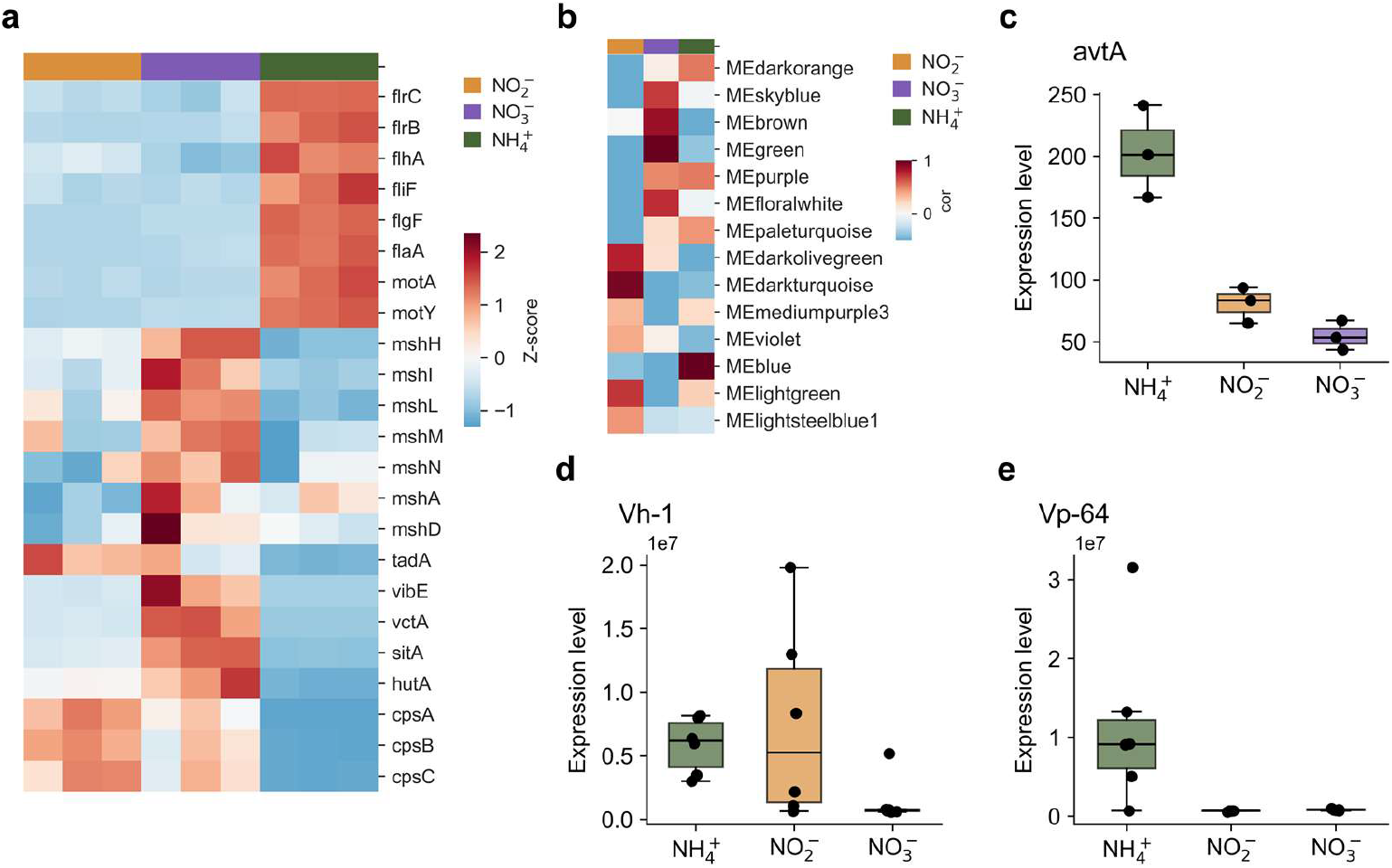
NH_4_^+^-induced transcriptional changes in Vh-1. **(a)** Heatmap showing the gene expression profiles under different nitrogen conditions. **(b)** Heatmap showing the correlation matrix between WGCNA modules and different cultivation conditions. **(c)** Expression level of *avtA*. **(d)** Alanine concentration in the culture medium of Vh-1. **(e)** Alanine concentration in the culture medium of Vp-64.

Given that flagellar biosynthesis represents an energetically costly cellular process, we further investigated metabolic changes associated with NH_4_^+^ utilization. Transcriptomic analysis revealed that *glnA* was highly expressed under both NH_4_^+^ and NO_2_^−^ conditions, indicating enhanced assimilation of ammonium-derived nitrogen. Moreover, *avtA* exhibited specific upregulation under NH_4_^+^ cultivation, suggesting that the abundant ammonium supply enhanced nitrogen flux toward amino acid biosynthesis and increased the potential for alanine production through glutamate-dependent transamination (Fig. 4c). Metabolomic analysis revealed significant accumulation of alanine in the Vh-1 culture medium under NH_4_^+^ cultivation (Fig. 4d). Similarly, alanine showed increased relative abundance in Vp-64 under NH_4_^+^ condition (Fig. 4e). These results suggest that the highly accessible ammonium source promotes rapid nitrogen assimilation, while excess nitrogen flux relative to biomass synthesis may cause an imbalance in carbon–nitrogen metabolism, leading to alanine accumulation and a potential metabolic overflow response.

Together, these findings indicate that NH_4_^+^ utilization drives a distinct physiological strategy characterized by enhanced motility and active nutrient acquisition, accompanied by metabolic overflow associated with excessive nitrogen assimilation.

### Nitrate availability induces a colonization-associated adaptation strategy in *V. harveyi*

Compared with NH_4_^+^ cultivation, NO_3_^−^ availability induced a distinct transcriptional program characterized by enhanced colonization-associated traits. Genes involved in surface attachment and extracellular matrix production, including capsular polysaccharide biosynthesis genes (*cps*) and the pilus genes (*msh*), exhibited elevated expression under NO_3_^−^ conditions, whereas flagellar-associated genes showed relatively reduced expression (Fig. 4a). This expression pattern contrasted with the NH_4_^+^ condition, where motility-related pathways were preferentially activated, suggesting a shift from motility-driven exploration toward a surface-associated lifestyle under nitrate availability. In addition to surface adaptation, genes involved in iron acquisition were strongly induced under NO_3_^−^ conditions. Key components of iron uptake systems, including the vibriobactin biosynthesis gene *vibE*, the vibriobactin receptor *vctA*, and iron transport-related genes such as *hutA* and *sitA*, were significantly upregulated (Fig. 4a).

Together, these transcriptional responses indicate that NO_3_^−^ availability promotes a colonization-associated adaptation strategy involving increased surface attachment, extracellular matrix production, and nutrient acquisition capacity in *V. harveyi*.

### Nitrite assimilation is coupled with redox stress adaptation in *V. harveyi*

Nitrite assimilation mediated by the *nirBD* system enables *V. harveyi* to satisfy its nitrogen demand. Meanwhile, this process may also reflect the simultaneous intracellular accumulation and detoxification of NO_2_^−^. The MEdarkturquoise module was identified as specifically associated with NO_2_^−^ cultivation in gene co-expression network, representing genes highly expressed under NO_2_^−^ condition (Fig. 4b). This module was primarily enriched in redox stress response pathways. The coordinated upregulation of oxygen-insensitive NAD(P)H nitroreductase, alkyl hydroperoxide reductase, cytochrome c peroxidase, FAD/SDR family oxidoreductases, and *OsmC* family proteins indicated activation of antioxidant and redox defense systems under NO_2_^−^ conditions.

Together, these responses indicate that NO_2_^−^ exposure represents a redox challenge for *V. harveyi*, leading to the activation of detoxification and antioxidant systems that support intracellular redox homeostasis.

## Discussions

### Differences in survival and stress tolerance strategies between *Vibrio* species

In this study, we compared the growth dynamics and population viability of two *Vibrio* species under conditions of limited carbon and nitrogen availability. Our results revealed substantial differences between the two species in inorganic nitrogen utilization strategies and long-term survival capacity.

For *V. harveyi*, rapid population expansion was accompanied by a relatively rapid decline in population viability, particularly under NH_4_^+^ condition. Although bacterial biomass continued to increase, population viability began to decrease before reaching the maximum cell density, suggesting that rapid growth may accelerate nutrient depletion and accumulation of metabolic by-products, thereby causing premature loss of cellular activity. In contrast, under NO_2_^−^ condition, *V. harveyi* exhibited a slower growth rate but continuously increased population viability, indicating that reduced growth rates may favor the maintenance of cellular physiological status. NO_2_^−^ is generally considered a potentially toxic nitrogen compound that can disrupt cellular redox homeostasis and induce oxidative stress, thereby restricting microbial growth. However, our results demonstrated that *V. harveyi* was capable of utilizing NO_2_^−^ as the sole inorganic nitrogen source to support continuous growth and induced the expression of assimilatory nitrite reduction-related genes, indicating a strong capacity for nitrite adaptation and utilization.

Compared with *V. harveyi*, *V. parahaemolyticus* exhibited a stronger ability to maintain population viability. Under NH_4_^+^ condition, *V. parahaemolyticus* not only achieved rapid proliferation but also maintained high population viability during long-term cultivation. Meanwhile, under NO_3_^−^ and NO_2_^−^ condition, although no obvious population expansion was observed, bacterial viability remained close to the initial inoculation level. These results indicate that although *V. parahaemolyticus* lacks the ability to efficiently utilize nitrate and nitrite for biomass production, it possesses strong long-term survival capacity. Since inorganic nitrogen availability may become a limiting factor during later cultivation stages, we speculate that the enhanced viability maintenance of *V. parahaemolyticus* may be associated with continued utilization of organic carbon sources, reduced metabolic activity, or enhanced starvation tolerance. Further investigation is required to elucidate the underlying mechanisms.

Overall, the two *Vibrio* species exhibited distinct ecological adaptation strategies. *V. harveyi* possesses a broader inorganic nitrogen utilization spectrum, enabling growth using NH_4_^+^, NO_3_^−^, and NO_2_^−^, but rapid proliferation is accompanied by accelerated loss of viability. In contrast, *V. parahaemolyticus* primarily relies on NH_4_^+^ for rapid growth but demonstrates stronger long-term survival capacity. These differences may reflect divergent ecological strategies employed by Vibrio species in oligotrophic environments.

### Heterotrophic nitrification suggests unexplored nitrogen metabolic potential in *Vibrio*

Heterotrophic nitrification refers to the process in which heterotrophic microorganisms utilize organic carbon sources for growth while oxidizing reduced nitrogen compounds into oxidized nitrogen forms, such as nitrite or nitrate. The conversion of ammonium to nitrate generally involves two sequential steps, namely ammonia oxidation and nitrite oxidation. These processes rely on key enzyme systems including *amo*, *hao*, and *nxr*, which have been reported in diverse microorganisms^24–26^. In this study, NO_3_^−^ production was detected during NH_4_^+^ cultivation of *V. harveyi.* Genome analysis further demonstrated that both *Vibrio* strains lacked *amo,hao* and *nxr*, indicating that their potential nitrification process differs from classical autotrophic nitrification and may rely on unidentified noncanonical metabolic mechanisms.

Given the transient accumulation of low levels of NO_2_^−^ during NH_4_^+^ cultivation and the subsequent detection of trace NO_3_^−^ during NO_2_^−^ cultivation, we hypothesized that nitrite oxidation may represent a potential intermediate step contributing to nitrate formation in *V. harveyi*. Recent studies have reported that the periplasmic nitrate reductase *napA* may exhibit catalytic activity for NO_2_^−^ oxidation to NO_3_^−^ in *Campylobacter jejuni* based on in vitro assays^23^. Interestingly, our transcriptomic analysis revealed that *napA* was significantly upregulated under NH_4_^+^ conditions in *V. harveyi*. Despite limited sequence similarity, the predicted structure of *napA* displayed a highly similar overall fold to *nxrA*, suggesting conservation of structural features within the molybdenum enzyme superfamily. Although this reverse catalytic activity of *napA* has not been experimentally validated in vivo, these findings suggest that enzymes traditionally assigned to specific nitrogen metabolic pathways may possess broader catalytic potential, providing new insights into the molecular mechanisms underlying heterotrophic nitrogen transformation.

Collectively, our findings suggest that homology-based functional annotation may not fully capture the complexity of nitrogen metabolism networks in *Vibrio*. Unidentified nitrogen metabolic enzymes, noncanonical catalytic mechanisms, and regulatory networks may remain undiscovered. Future integration of proteomics, metabolic flux analysis, and biochemical validation will be essential for elucidating the molecular mechanisms underlying heterotrophic nitrification and inorganic nitrogen utilization in *Vibrio*.

### Differential nitrogen sources reshape life-history-associated transcriptional programs in *Vibrio*

Although *V. parahaemolyticus* Vp-64 exhibited limited growth capacity when utilizing NO_3_^−^ and NO_2_^−^, its genome still harbored multiple nitrogen metabolism-related genes. The manifestation of nitrogen metabolism-related phenotypes may depend on multiple factors, including transcriptional regulation, compatibility of electron transport systems, cofactor availability, and environmental conditions, such as carbon source composition and oxygen availability. For example, the presence of glucose as the primary carbon source and non-anaerobic cultivation conditions may influence the activation of nitrogen metabolic pathways. Therefore, the limited utilization of NO_3_^−^ and NO_2_^−^ by *V. parahaemolyticus* may reflect insufficient activation of these pathways under the tested conditions rather than the absence of corresponding genetic modules.

Our results demonstrate that *V. harveyi* Vh-1 possesses the capacity to assimilate and utilize NO_2_^−^ as a nitrogen source. However, compared with NH_4_^+^ and NO_3_^−^ conditions, the *nirBD* genes exhibited the lowest expression levels under NO_2_^−^ cultivation. This finding indicates that *nirBD* expression is not solely induced by nitrite availability but is also regulated by the overall cellular nitrogen metabolic status. Under NO_3_^−^ conditions, cells must undergo sequential nitrate reduction, in which nitrate is first reduced to nitrite and subsequently converted into ammonium^1,9^. Therefore, the elevated expression of the *nirBD* genes may facilitate the rapid assimilation of nitrite intermediates generated during nitrate reduction, preventing their accumulation and ensuring efficient nitrogen flux toward biosynthetic pathways. In contrast, although exogenous nitrite can directly serve as a substrate for the *nirBD* pathway under NO_2_^−^ conditions, the relatively lower expression of *nirBD* may reflect metabolic adaptation strategies associated with limited nitrite assimilation capacity or nitrite-induced stress. This hypothesis is further supported by the elevated expression of multiple antioxidant-related enzymes under NO_2_^−^ cultivation.

*V. harveyi* was capable of growth under both NH_4_^+^ and NO_3_^−^ conditions and exhibited distinct nitrogen metabolic patterns. Together with the observation of nitrate production during NH_4_^+^ cultivation, these results suggest that *V. harveyi* may adapt to dynamic changes in environmental nitrogen availability through noncanonical heterotrophic nitrification and nitrate assimilation pathways. Transcriptomic analysis further demonstrated that inorganic nitrogen sources not only altered nitrogen metabolism-related gene expression but also induced systematic remodeling of multiple virulence-associated functional modules, including flagella, pili, and siderophore systems, suggesting that nitrogen availability may drive distinct adaptation strategies.

Flagella are major structures responsible for *Vibrio* motility and chemotaxis, allowing cells to actively explore favorable niches and initiate host interactions^27,28^. Pili and capsular polysaccharides mediate surface attachment, biofilm development, and persistent colonization, representing key determinants of the transition from planktonic to surface-associated lifestyles^29–31^. Siderophores promote iron acquisition in iron-limited environments and support bacterial proliferation, ecological competition, and pathogenicity^32,33^. Therefore, these functional modules are not only involved in host infection but also represent important components underlying transitions among free-living, environmental colonization, and host-associated lifestyles. In our study, flagellar-associated genes were generally upregulated under NH_4_^+^ cultivation, suggesting enhanced motility-related capacity. NH_4_^+^ represents a readily accessible nitrogen source that can be directly incorporated into central nitrogen assimilation pathways. Although flagellar synthesis and operation are energy-intensive processes, efficient ammonium assimilation may exceed the capacity for biomass production under nitrogen-rich conditions, resulting in an imbalance between carbon flux and nitrogen utilization. Consistently, alanine significantly accumulated in the culture medium of both *Vibrio* strains under NH_4_^+^ cultivation. This extracellular alanine accumulation may represent a metabolic overflow response during ammonium utilization, in which excess metabolic intermediates are redirected toward amino acid production and release to maintain intracellular metabolic homeostasis^34,35^.

In contrast, under NO_3_^−^ conditions, *V. harveyi* exhibited coordinated induction of pilus-and capsular polysaccharide-associated genes, together with siderophore biosynthesis and transport systems. On one hand, the elevated expression of pilus and capsule-related gene suggests that cells may enhance surface attachment, biofilm formation, and environmental colonization-associated capabilities, reflecting an adaptive shift from a free-living lifestyle toward an attachment-oriented lifestyle. On the other hand, activation of siderophore-associated pathways indicates that *V. harveyi* may improve its environmental resource competition through enhanced iron acquisition, thereby supporting long-term survival and ecological adaptation under nutrient-limited conditions. Collectively, the surface structural remodeling and resource acquisition strategies induced under NO_3_^−^ conditions may represent an environmental persistence-oriented adaptation strategy distinct from the high-motility response observed under NH_4_^+^ conditions.

Taken together, our findings suggest that inorganic nitrogen sources function not only as metabolic substrates but also as important environmental signals regulating life-history strategies, metabolic allocation, and potential virulence-associated traits in *V. harveyi*. These results provide new insights into how *Vibrio* species integrate environmental nitrogen availability with ecological adaptation and pathogenic potential in aquatic ecosystems and host-associated environments.

## Methods

### Bacterial strains

The *V. harveyi* and *V. parahaemolyticus* strains used in this study were isolated from muscle tissues of diseased crab collected from shrimp farms in southern China. The bacterial strains were preserved in glycerol stocks at −80°C. Genome assemblies and annotations were obtained from NCBI RefSeq, with accession number PRJNA749085 and PRJNA716109.

### M9 medium preparation

The M9 medium was prepared as follows. The phosphate buffer system consisted of 17.1 g/L Na_2_HPO_4_·12H_2_O and 3.0 g/L KH_2_PO_4_. Glucose was supplied as the sole carbon source at a final concentration of 2 g/L. Sodium chloride was added at 10 g/L. Magnesium sulfate and calcium chloride were supplemented at final concentrations of 1 mM each. Three inorganic nitrogen sources, including NaNO_3_, NH_4_Cl, and NaNO_2_, were separately added to provide an equivalent amount of nitrogen, corresponding to 38 mg/L of elemental nitrogen (N).

### Bacterial activation, inoculation, and cultivation

Frozen stocks of *V. harveyi* and *V. parahaemolyticus* were directly inoculated into LB liquid medium for cultivation. Bacterial cultures were pre-grown at 28°C with shaking at 200 rpm for 24 h. After cultivation, cells were harvested by centrifugation at 5000 × g for 5 min, and the supernatant was discarded. The bacterial pellets were washed three times with sterile saline (10 g/L NaCl) to remove residual nutrients from the LB medium.

The bacterial concentration was estimated by measuring OD_600_ using a microplate reader. Based on the measured OD_600_ values, the required inoculum volume was calculated to achieve an initial bacterial density of 1 × 10^5^ CFUs/mL. Cultivation was performed in 50 mL centrifuge tubes containing 5 mL M9 medium supplemented with one of the three inorganic nitrogen sources. After inoculation, cultures were incubated at 30°C with shaking at 200 rpm. After 12 h of cultivation, OD_600_ values were measured every 2 h. Viable bacterial populations were quantified by plating serially diluted cultures on TCBS agar plates during cultivation.

### Detection of three inorganic nitrogen forms in culture medium

The concentrations of three inorganic nitrogen forms in the culture medium were quantified by Guangzhou Dinghai Technology Co., Ltd. (Guangzhou, China). NH_4_^+^-N was quantified using the Nessler reagent spectrophotometric method, and NO_2_^−^-N was measured using the diazotization-coupling spectrophotometric method. NO_3_^−^-N was determined using ultraviolet spectrophotometry, with sulfamic acid treatment applied to remove interference caused by residual nitrite in the samples.

### Protein structure prediction and structural similarity analysis

Protein sequences of *V. harveyi* were obtained from RefSeq genome annotations. Proteins with lengths ≤1500 amino acids were selected for tertiary structure prediction and downstream structural analysis. Reference *nxrA* sequences were collected from experimentally characterized *nxrA* structures and homologous clustered sequences retrieved from UniRef^36^. For UniRef-derived sequences, only proteins longer than 800 amino acids were considered potential *nxrA* homologs and included in subsequent structural comparisons (Supplementary Materials).

Protein tertiary structures were predicted using ColabFold (v1.5.5), which implements the AlphaFold2 framework^37^. For each protein sequence, three AlphaFold2 parameter sets (models 1–3) were evaluated with three recycling iterations to improve prediction accuracy. Template information was enabled during prediction, and the resulting models were subjected to Amber-based energy minimization. A fixed random seed of 42 was applied to ensure reproducibility. Structure prediction was terminated early when the predicted model confidence score exceeded 70, and the model with the highest confidence score was selected for subsequent structural analyses.

Structural similarity analysis was performed using Foldseek (v10) by comparing predicted *V. harveyi* protein structures with reference *nxrA* structures^38^. Candidate proteins with a TM-score above 0.5 and coverage above 0.7 were selected as potential structural homologs of *nxrA*.

### Transcriptome analysis

Raw sequencing reads were subjected to quality assessment using Fastp (v0.20.1), including evaluation of sequencing quality distribution, adapter contamination, and nucleotide composition bias^39^. The reference genome sequences and corresponding annotation files of *V. harveyi* were downloaded from NCBI RefSeq and used to construct genome indices. Gene expression levels were quantified using Salmon (v 1.10.1)^40^.

Differential expression analysis was performed using DESeq2 (v1.34.0), including normalization, dispersion estimation, and negative binomial model fitting^41^. Differentially expressed genes (DEGs) were identified using the criteria of an absolute fold change ≥3 and an adjusted p-value <0.05. Genes with more than 10 mapped short reads in at least two samples were retained for downstream analysis. Gene Ontology (GO) enrichment analysis of DEGs was performed using the ClusterProfiler (v4.2.2)^42^.

Gene co-expression network analysis was performed using WGCNA (version 1.72-5)^43^. The soft-thresholding power of 12 was selected, at which the scale-free topology fit index reached 0.8, indicating a scale-free network structure.

### Metabolite extraction and sample preparation

Culture samples were centrifuged at 5,000 × *g* for 4 min, and 1 mL of the supernatant was collected and filtered through a 0.22 μm membrane. An appropriate volume of filtrate was transferred into a 2 mL centrifuge tube, followed by the addition of 300 μL methanol/acetonitrile (V/V=2:1) extraction solution. Samples were vortexed for 1 min and ultrasonicated in an ice bath for 10 min. After centrifugation at 12,000 rpm for 10 min at 4°C, 300 μL of the supernatant was transferred into a new 2 mL centrifuge tube and dried under vacuum concentration. The dried extracts were reconstituted in 150 μL methanol/water solution (V/V=1:4) containing 2-chloro-L-phenylalanine as an internal standard. After centrifugation, the supernatants were filtered through a 0.22 μm membrane, transferred into injection vials, and subjected to LC–MS analysis. LC–MS-based metabolomic analysis was conducted with technical assistance from Tgene Biotech (Shanghai) Co., Ltd.

### Metabolomics data processing and analysis

Raw mass spectrometry files were converted using the Proteowizard (v3.0.8789)^44^. Peak detection, filtering, and alignment were performed using the XCMS (v3.12.0), generating a quantitative metabolite matrix^45^. The parameters were set as follows: *bw* = 2, *ppm* = 15, *peakwidth* = c (5, 30), *mzwid* = 0.015, *mzdiff* = 0.01, and *method* = "centWave". Support vector regression-based correction using quality control samples was applied to reduce systematic errors. Metabolites with relative standard deviation values >30% in QC samples were removed during quality control procedures before downstream analysis.

Metabolite annotation was performed by matching experimental MS/MS spectra against multiple databases, including HMDB^46^, MassBank^47^, LipidMaps^48^, mzCloud, KEGG, and the in-house standard library established by Tgene Biotech (Shanghai) Co., Ltd. The mass tolerance was set to <30 ppm. Metabolite identification was performed based on precursor ion mass-to-charge ratios from MS1 spectra, mass deviation, and adduct information for molecular formula prediction, followed by MS/MS fragment ion matching against database records for secondary metabolite identification.

Statistical analysis was performed using the ropls (v.1.44.0)^49^. *P* values were calculated for statistical significance testing, orthogonal partial least squares discriminant analysis was performed for dimensionality reduction and sample separation, and variable importance in projection scores and fold changes were calculated to evaluate differential metabolites between groups.

### Quantitative real-time PCR

Quantitative real-time PCR was performed using a Bio-Rad CFX96 Real-Time PCR Detection System (Bio-Rad, CA, USA). Each qRT-PCR reaction was performed in a final volume of 10 μL, containing 0.8 μL cDNA template, 5 μL TB Green Premix Ex Taq II (Tli RNaseH Plus) (2×), 0.4 μL forward primer, 0.4 μL reverse primer, and 3.4 μL RNase-free ddH_2_O. The amplification program was as follows: initial denaturation at 95°C for 30 s, followed by 40 cycles of amplification consisting of 95°C for 5 s and 60°C for 30 s. Primer sequences used for qRT-PCR validation are listed in S. Table 2.

## Data Availability

The raw matrix and other supplementary files have been uploaded to https://zenodo.org/records/21350600. Genome assemblies can be obtained from NCBI with accession number PRJNA749085 and PRJNA716109.

## Author Contributions

Study conceptualization by L.G. and X.X. The experiments were conducted under the supervision of H.R. and L.G. Data analyses were performed by X.X. with support from H.R. The manuscript was written by X.X. with substantive contributions from L.G. Funding acquisition by L.G.

## Acknowledgement

This work was supported by the fund of the Zhuhai Social Development Science and Technology Project (2420004000111), Modern Agriculture Industry Technology Innovation Teams, Department of Agriculture and Rural Affairs of Guangdong Province (2024CXTD26), Plan Project of Science and Technology of Heyuan (grant no. Heke2023001 and Heke2021039). We extend our sincere thanks to Zhaolin lv, Canming Yang, Dr. Dandan Huang for their guidance, and helpful discussions.

